# PDE3A2 modRNA Fine-Tunes Nuclear cAMP Microdomains and Reverses Pathological Cardiac Hypertrophy

**DOI:** 10.1101/2025.09.30.679675

**Authors:** Wenjing Xiang, Mingwei Lian, Qiuyun Guan, Xingsheng Sun, Supeng Li, Haocheng Lu, Gangjian Qin, Yang Kevin Xiang, Haibo Ni, Ying Wang

**Author notes:** Contributed equally (Wenjing Xiang, Mingwei Lian, Qiuyun Guan). **Corresponding authors:** Haibo Ni, PhD,. Ying Wang, PhD, No. 1088, Xueyuan Avenue, Nanshan District, Shenzhen City, Guangdong Province, 518055, China,.

## Abstract

Pathological cardiac hypertrophy (PCH) is a precursor to heart failure, driven in part by dysregulated nuclear-localized cAMP (NLS-cAMP) signaling. Phosphodiesterase 3A2 (PDE3A2) is a key regulator of this nuclear cAMP microdomain, yet its selective modulation remains challenging. Here, we developed PDE3A2-modified RNA (PDE3A2-modRNA) to restore nuclear PDE3A2 levels and tested its efficacy in an angiotensin II-induced PCH model. Subcellular FRET imaging revealed that PCH hearts exhibit hyperactive NLS-cAMP due to PDE3A2 depletion. PDE3A2-modRNA selectively reduced NLS-cAMP without altering cytosolic cAMP, demonstrating precise microdomain regulation. *In vivo*, PDE3A2-modRNA improved cardiac function, attenuated hypertrophy and fibrosis, and shifted transcriptional programs toward physiological remodeling. Single-nucleus transcriptomics and immune profiling further revealed reduced oxidative stress and an improved cardiac microenvironment. These findings highlight PDE3A2-modRNA as a novel gene therapy that selectively restores nuclear cAMP homeostasis, reversing PCH-promoting transcriptional programs while preserving physiological cAMP signaling in cytosolic microdomains.

## Introduction

Pathological cardiac hypertrophy is a maladaptive response of heart wherein cardiomyocytes undergo enlargement upon various hormonal and mechanic stimuli^1^. The development and progression of PCH are driven by a rewired gene transcription program that encodes the structural and functional signatures in PCH. PCH often progresses to heart failure, causing significant mortality worldwide. Thus, there is an urgent clinical need to treat PCH to lower the risk of heart failure. However, effective therapeutics for PCH remain elusive^2,3^.

Cyclic nucleotide signaling emerges as a key regulator of gene transcription program that is involved in pathological conditions, including PCH^4^. Recent studies have demonstrated that cAMP regulates gene expression by modulating histone transcription^5,6^. Involvement of cAMP in PCH was also supported by that overexpressing inhibitory peptide of PKA downstream signaling of (the major downstream effector of cAMP) could ameliorate PCH in mice^7^. Therefore, cAMP signaling pathway is a potential effective target for PCH. Our previous studies have demonstrated that cAMP is organized into discrete subcellular microdomains that enables the precise spatiotemporal control of intracellular signaling and diverse cell function^8,9^, including cardiac excitation-contraction coupling^10–13^. Therefore, it is crucial to tease out the dysregulated cAMP microdomains that cause pathology from those that are responsible for maintaining cardiac function for development of cAMP-targeted treatment of PCH. Importantly, this cAMP microdomain-biased approach may leverage the isoform specific localization of phosphodiesterases (PDEs), which selectively fine-tune the discrete cAMP microdomain by degrading local cAMP in specific subcellular locations ^13^. PDE3 inhibitors are clinically used for acute treatment of heart failure but cause increased mortality with its long-term use^14^. Among two isoforms of PDE3A, PDE3A2 but not PDE3A1 has been shown to localize to the nuclei of neonatal cardiomyocytes and PDE3 inhibition leads to the hypertrophy of neonatal cardiomyocytes^6^, positioning PDE3A2 as a promising therapeutic target for PCH by regulating nuclear cAMP microdomain. However, its *in vivo* role in PCH has yet to be fully elucidated.

While targeting a specific PDE isoform with clinically available drugs is highly challenging, gene therapy emerges as an attractive alternative therapeutic approach. Modified RNA (modRNA)-based gene therapy offers significant advantages, enabling the delivery of specific proteins with high efficiency, transient expression, enhanced safety, and non-immunogenicity^15,16^. Ongoing clinical and preclinical trials of modRNA focusing on ischemic heart repair by targeting myocyte proliferation and angiogenesis^15^. Leveraging modRNA to target specific PDE isoforms holds potential for precisely modulating pro-hypertrophic gene transcription. However, no modRNA-based therapies targeting cardiac hypertrophy have been reported.

In this study, using Förster resonance energy transfer (FRET) biosensors, we uncovered a hyperactive cAMP signaling in the nuclei but not at the plasma membrane (PM) of cardiomyocytes from PCH mice. The aberrant NLS-cAMP was associated with reduced PDE3A2 expression. Using PDE3A2-modified RNA (PDE3A2-modRNA) technology, we found that PDE3A2-modRNA treatment suppressed NLS-cAMP of cardiomyocytes and attenuated cardiac remodeling and improved cardiac function of PCH mice. Single-cell nuclei RNA sequencing unraveled that PDE3A2-modRNA promoted physiological hypertrophy-associated signaling while suppressing those associated with PCH. PDE3A2-modRNA attenuated injury-responding inflammatory cardiomyocyte functional trajectory. In addition, PDE3A2-modRNA rebuilt microenvironment by attenuating complement activation in both cardiomyocytes and non-cardiomyocytes of PCH hearts. Our study presents the first modRNA-based gene therapy targeting PDE isozymes as a promising and effective therapeutic approach for PCH.

## Results

### Hypertrophic hearts exhibited an aberrant increase in nuclear cAMP levels with reduced expression of PDE3A

PCH is characterized by dramatic alterations in gene transcription programs, which are tightly controlled by cAMP-PKA signaling^3,7^. To investigate the role of nuclear localization sequence (NLS)-cAMP in PCH, we employed a mouse model^26–28^ of PCH induced by angiotensin II (Ang II, 1000 ng/kg/min, 14 days) infusion using osmotic minipumps. Two-week infusion of Ang II significantly increased heart size and heart weight to tibia length ratio (**Figure 1A-B**), indicating the development of PCH. Also, wheat germ agglutinin (WGA) staining of mouse hearts revealed an increase of cardiomyocyte size in PCH vs control (ctrl) groups (**Figure 1C**). To dissect the subcellular-localized cAMP-PKA signal changes induced by PCH, we isolated cardiomyocytes from mouse hearts and infected them with adenovirus to express subcellularly-localized FRET-based PKA biosensor (AKAR3), targeted to either the plasma membrane (PM) or the nucleus (**Figure 1D-I** and **Online Figure 1A-C**). FRET and fluorescence imaging confirmed that NLS-AKAR3 was exclusively expressed in the nuclei of myocytes (**Figure 1G**) and 293A cells (**Online Figure 1B-C**), while PM-AKAR3 localized to the cellular surface of myocytes and 293A cells (**Figure 1G** and **Online Figure 1B-C**). Upon norepinephrine (NE, 100 nM) stimulation, we observed distinct PCH-dependent alteration in subcellular PKA activity. PKA activity at PM (PM-PKA) was markedly decreased in PCH vs ctrl (**Figure 1E-F**), which could be due to the desensitization of adrenergic receptors in PCH^4,29^. In contrast, nuclear PKA activity (NLS-PKA) hyperactivated in PCH group (**Figure 1H-I**), suggesting an aberrant increase in NLS-cAMP.

**Figure 1.**
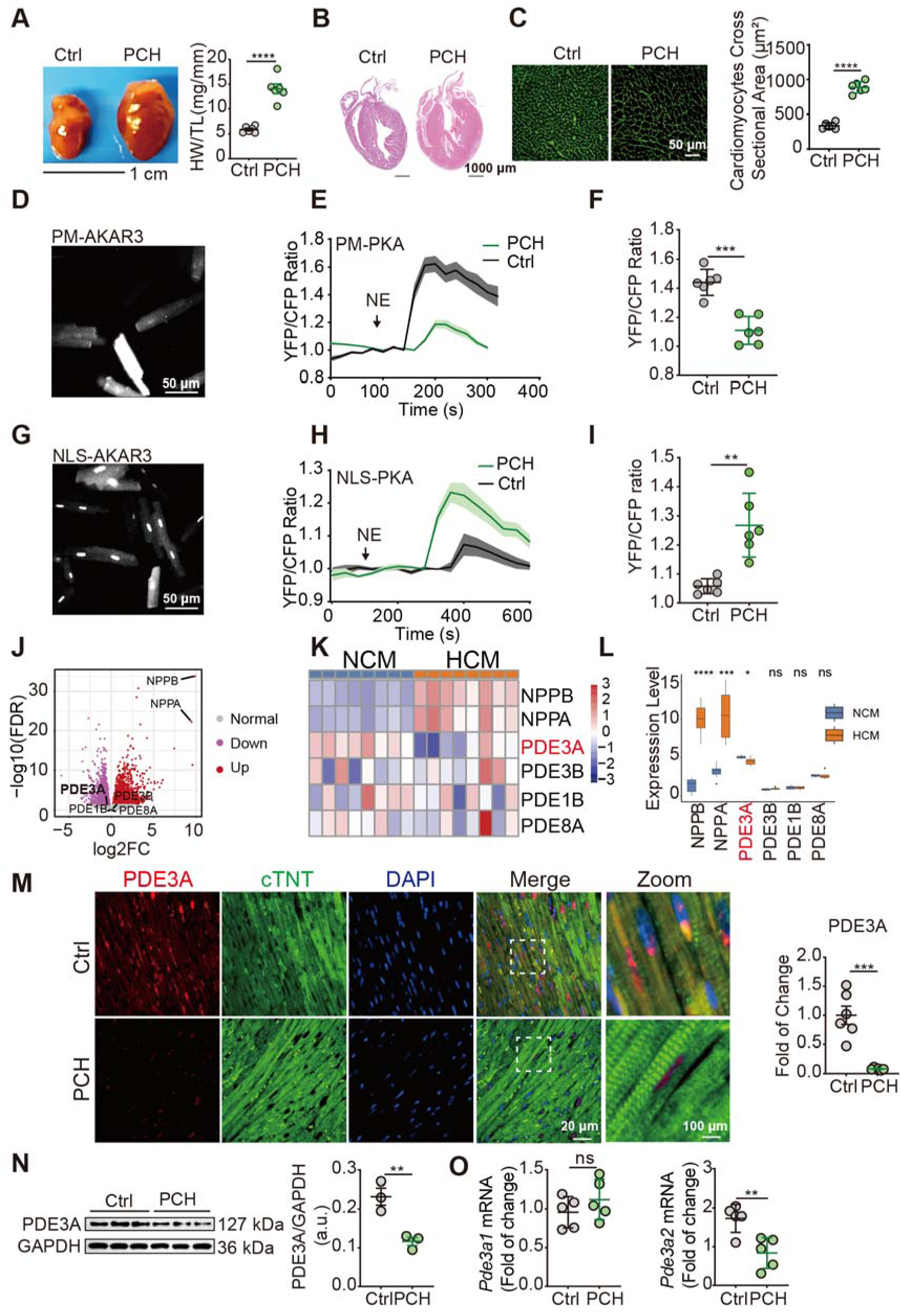
PCH is associated with overactivated nuclear-localized cAMP-PKA and down-regulated PDE3A2. (**A-B**) Representative heart imaging and heart weight to tibia length ratio (mg/mm) in healthy control (ctrl) and angiotensin II (Ang II)-induced PCH mice. (**C**) Representative WGA staining and quantification of cardiomyocyte area in Ctrl and PCH mouse hearts. (**D-F**) Cardiomyocytes isolated from Ctrl and PCH mice expressing plasma membrane (PM)-localized or nuclear-localized (NLS) PKA reporter (AKAR) were stimulated with norepinephrine (NE). Representative imaging and typical curve of PM-PKA in response to NE. Summary of PM-PKA response in Ctrl and PCH cardiomyocytes. (**G-I**) Representative imaging and typical curve of NLS-PKA in response to NE. Summary of NLS-PKA response in Ctrl and PCH cardiomyocytes. (**J**) Volcano plot of up and down regulated genes in human hypertrophic cardiomyopathy (HCM, dataset accession number: GSE133054). (**K**) Heatmap of pro-hypertrophic genes and PDEs in HCM and non-cardiomyopathy control (NCM). (**L**) Boxplot quantifications of gene expression levels in HCM and NCM. (**M-N**) Quantification of PDE3A expression in Ctrl and PCH groups by immune fluorescence staining (**M**) and immunoblot (**N**). (**O**) The mRNA levels of *Pde3a1* and *Pde3a2* in Ctrl and PCH groups. Ang II (1000 ng/kg/min, 14 days) was infused to mouse by osmotic minipump to induce PCH. Data were shown as mean ± SD. n = 6 mice for each group. *P* values were obtained by unpaired student *t*-test. \**p* < 0.05, \*\**p* < 0.01, \*\*\**p* < 0.001, \*\*\*\**p* < 0.0001.

Phosphodiesterases (PDEs) with different subcellular localizations regulate specific cAMP pools^13^, and dysregulation of PDE-mediated cAMP-PKA signaling contributes to PCH development^4^. To identify the PDE responsible for the excessive NLS-cAMP in PCH, we re-analyzed published RNA-seq data (Accession number: GSE133054) from patients with or without hypertrophic cardiomyopathy (HCM)^2^ (**Figure 1J-L**). Bulk RNA sequencing was performed on left ventricular samples from control and hypertrophic samples. Compared to non-cardiomyopathy (NCM) samples, the HCM groups show marked upregulation of NPPA and NPPB, fetal gene markers of hypertrophy. Among PDE isoforms, we found that PDE3A, but not other PDEs (e.g., PDE3B, PDE1B, and PDE8A), was decreased in HCM (**Figure 1 J-L**). To test whether PDE3A expression was similarly altered in our mouse PCH model, we performed immunostaining and western blot of mouse heart tissue (**Figure 1M-N**), which confirmed a reduction in PDE3A levels in PCH hearts. Interestingly, PDE3A was partially localized in the nuclei of cardiomyocytes (**Figure 1M**), consistent with previous observation in isolated cardiomyocytes^6^. Further qPCR analysis revealed that the reduction was specific to the *Pde3a2* isoform, with no significant changes in *Pde3a1* (**Figure 1O**). A previous study demonstrated that PDE3A2 regulates NLS-cAMP microdomain, and global inhibition of PDE3 promotes cardiomyocyte hypertrophy. However, the specific role of PDE3A2 *in vivo* remains to be fully elucidated^6^. Taken together, these findings suggest a link between hyperactive NLS-cAMP and diminished PDE3A2 expression, pointing to PDE3A2 as a potential therapeutic target for PCH.

### PDE3A2-modRNA abolished cAMP signal at nucleus without impacting cytosolic cAMP signaling

ModRNA technology offers the potential to selectively target specific PDE3A isoforms, a capability not achievable with current PDE3A-specific drugs. To explore this, we designed a modRNA encoding mouse PDE3A2 (**Figure 2A**). The open reading frame of PDE3A2 was cloned to pVAX1(+) for the high-yield RNA *in vitro* transcription and subsequently encapsulated in lipid microparticles (LNPs) (**Figure 2A**). Using this LNP-modRNA platform, we also generated LNP-encapsulated EGFP-modRNA. The encapsulated modRNAs had an average size of approximately 100 nm and exhibited high encapsulation efficiency (**Online Figure 2A**). Cell viability assays confirmed that both LNP and LNP-encapsulated EGFP-modRNA were safe to for 293A cells at concentrations of 1 μg, 5 μg and 20 μg/mL (**Online Figure 2B-C**), demonstrating the biosafety of our LNP-encapsulated modRNA. Furthermore, modRNA-PDE3A2 induced robust PDE3A2 protein expression in 293A cells, with increases ranging from 5-to 150-fold (**Online Figure 3A-C**). Notably, PDE3A2-modRNA-expressed proteins were predominantly located to the nucleus (**Online Figure 3B-C**). To examine if the delivered PDE3A2-modRNA could selectively modulate local cAMP-PKA signaling, we performed FRET imaging to compare nuclear and cytosolic PKA signals (**Figure 2B** and **2D**) using subcellular-localized FRET-based PKA biosensor^30^. PDE3A2-modRNA treatment prevented NLS-PKA activation upon stimulation of adenyl cyclase by forskolin (FSK, **Figure 2B**, **C**, **F**). This inhibition was reversed by IBMX, which rescued the NLS-PKA responses, regardless of PDE3A2-modRNA’s presence (**Figure 2B**, **C**, **F**). Conversely, PDE3A2-modRNA did not affect the cytosolic PKA activity following adenyl cyclase activation (**Figure 2D**, **2E** and **2F**), as measured using a nuclear export sequence (NES)-anchored PKA FRET biosensor (**Figure 2D**, **2E** and **2F**; **Online Figure 1B-C**). Under combined treatment with FSK and IBMX, both NLS-PKA and NES-PKA activity were comparable between Ctrl and PDE3A2-modRNA groups (**Figure 2F**). In summary, PDE3A2-modRNA selectively suppresses nuclear cAMP-PKA signaling without affecting cAMP signaling in other subcellular compartments.

**Figure 2.**
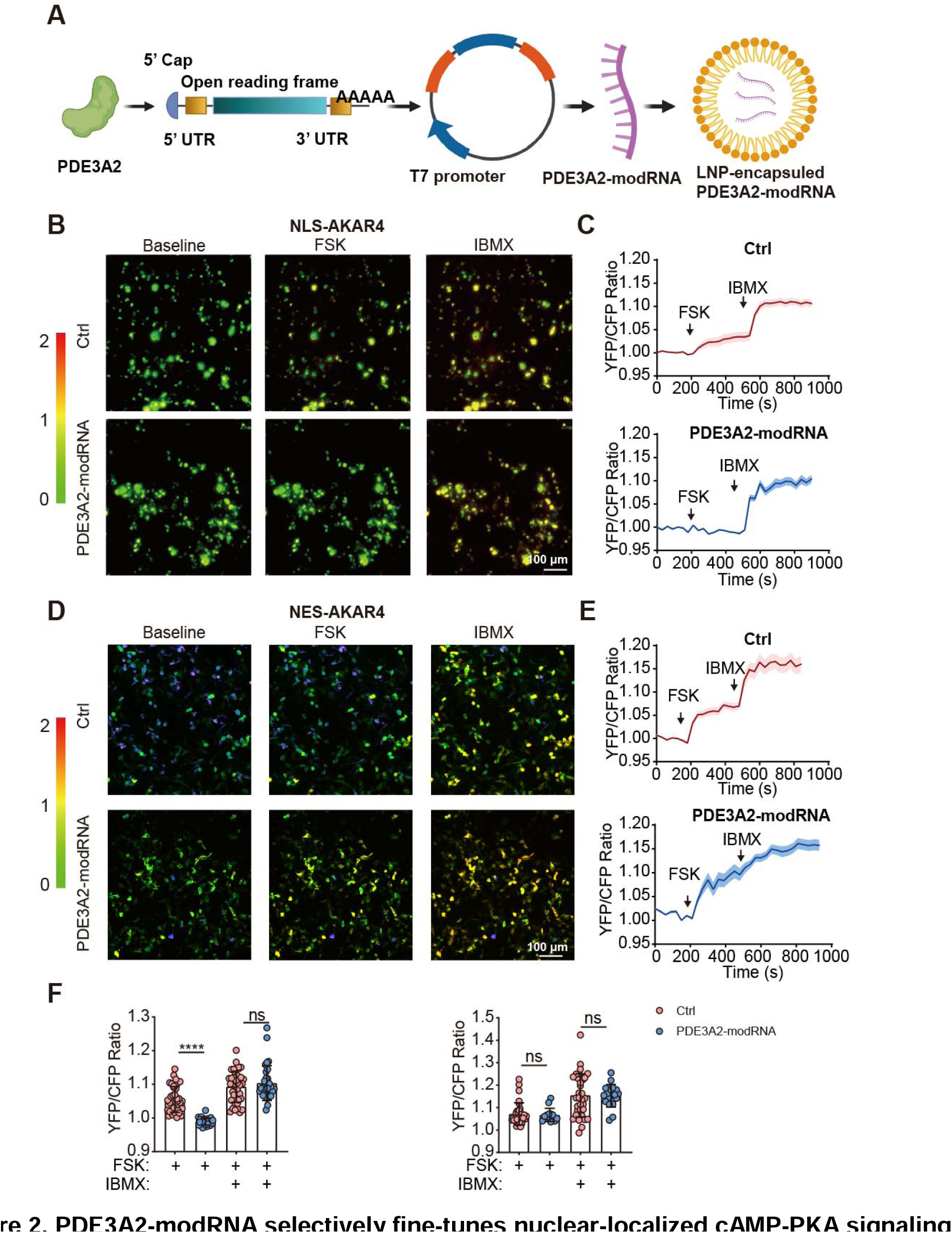
PDE3A2-modRNA selectively fine-tunes nuclear-localized cAMP-PKA signaling microdomain. **(A)** Pipeline for constructing mouse PDE3A2-modRNA with His-tag. **(B-C)** Representative FRET ratio images and typical curves of NLS-PKA response to the sequential stimulation of forskolin (FSK) or 3-Isobutyl-1-methylxanthine (IBMX) with or without PDE3A2-modRNA. (**D-E**) Representative FRET ratio images and typical curves of NES-PKA response to the sequential stimulation of FSK and IBMX. (**F**) Scatterplot summarizing NLS-PKA or NES-PKA FRET responses to FSK and IBMX. Pseudo-color FRET ratio images generated by YFP/CFP normalized to baseline. n = > 20 cells per conditions. FSK = forskolin, 10 μmol/L; IBMX = 3-Isobutyl-1-methylxanthine, 100 μmol/L. Data were shown as mean ± SD. *P* values were obtained by one-way ANOVA followed by Tukey’s post-hoc test. \*\*\*\**p* < 0.0001.

### PDE3A2-modRNA alleviated cardiac hypertrophic remodeling in Ang II-infused mice

We compared the efficiency of modRNA delivery through local cardiac injection and intravenous (*i.v.*) administration of EGFP-modRNA in mice hearts (**Online Figure 4**). Both intramyocardial injection and *i.v.* administration of 20 μg EGFP-modRNA resulted in substantial increase in EGFP expression in mouse hearts (**Online Figure 4A**, **B**) after 3 days. Meanwhile, no changes were detected in mouse systolic and diastolic function 3 days after the *i.v.* administration of EGFP-modRNA in healthy mice (**Online Figure 4C**, **D**). To minimize additional injury to the mouse heart caused by intramyocardial injection and enhance practicality, we chose *i.v.* administration for modRNA delivery in subsequent experiments. Additionally, heart morphology and Masson’s trichrome staining showed that EGFP-modRNA delivery did not affect the normal heart function and structure (**Online Figure 5A**, **B**). We then measured the biodistribution of *i.v.* administrated Fluc-modRNA to assess the efficiency of our modRNA-delivery to distinct organ. Consistent with previous study, our result suggests systemic *i.v.* administration induced protein expression in various organ, including the heart. (**Online Figure 6**). Together, these findings demonstrated that LNP-encapsulated modRNA injection efficiently deliver mRNA to the heart without significantly affecting the cardiac function.

Western blot and immunostaining confirmed the treatment of LNP-encapsuled PDE3A2-modRNA caused significantly increased PDE3A2 protein levels (by 6-7-fold) in cultured cardiomyocytes (**Figure 3A**). Of note, a majority of PDE3A2 was localized to the nuclei of cardiomyocytes (**Figure 3B**). Next, we sought to assess effects of PDE3A2-modRNA treatment on hearts from Ang II-infused mice. Ang II was infused for 14 days, followed by intravenous injections of PDE3A2-modRNA (20 μg, at day-15, 17, and 19, as shown in **Figure 3C**). PDE3A2-modRNA administration attenuated the increase in heart size, height weight (**Figure 3D**, **E**), and heart weight-to-tibia length ratio (**Figure 3E**), while also increasing the expression level of PDE3A in the myocardium (**Figure 3F**). Additionally, PDE3A2-modRNA reduced cardiomyocyte area in Ang II-infused mouse hearts (**Figure 3J**, **K**). Consequently, PDE3A2-modRNA normalized PCH mouse cardiac function, reduced the thickness of left ventricular posterior wall (**Figure 3G**, **H** and **Online Table 1-2**), and alleviated cardiac fibrosis (**Figure 3I**). Together, our data supports modRNA-mediated inhibition of nuclear-cAMP microdomains as a therapeutic strategy for PCH.

**Figure 3.**
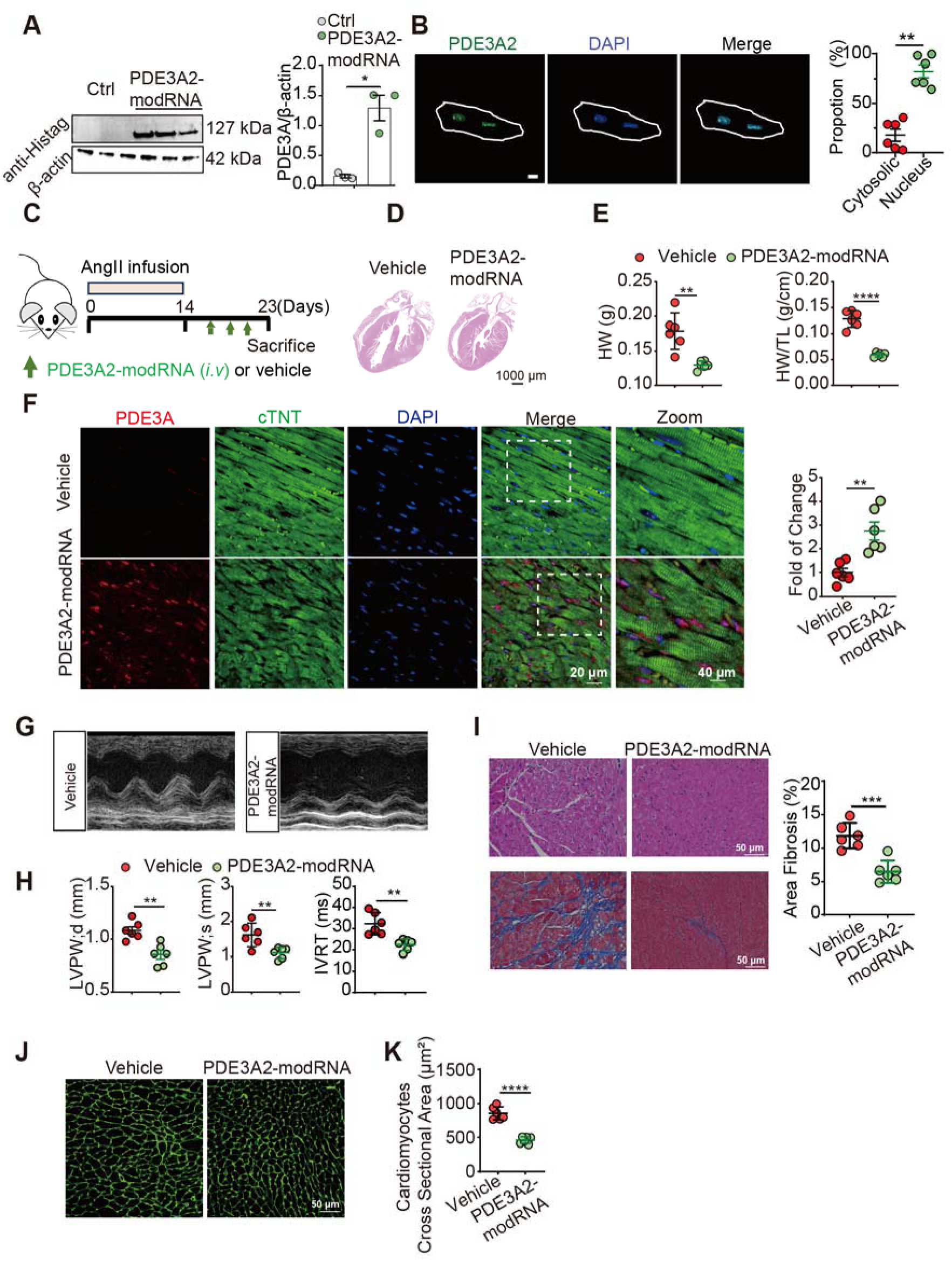
LNP delivery of PDE3A2-modRNA restores PDE3A2 expression level and alleviates mouse PCH. (**A**) PDE3A2 expression in the absence or presence of LNP-delivered PDE3A2-mRNA (30 ng/mol, 12 h) in isolated adult mouse ventricle cardiomyocytes (AVMs). (**B**) Distribution of PDE3A2 in AVMs. AVMs were stained with anti-His-tag antibody. (**C**) LNP vehicle control (Vehicle) or LNP PDE3A2-modRNA (PDE3A2) was *i.v.* administrated to mice after Ang II-infusion. PDE3A2-modRNA was given every two days three times as illustrated. (**D**) Exemplary images of mouse heart after treatment with Vehicle or PDE3A2-modRNA. (**E**) Effects of PDE3A2-mRNA on heart weight and heart weight to tibia length ratio. (**F**) Representative images and quantification of PDE3A in mouse hearts. Hearts were stained with anti-PDE3A antibody. (**G**-**H**) Echocardiography and measurements of the left ventricle thickness in vehicle or PDE3A2-modRNA treated mice. (**I**) Hematoxylin and Eosin (H&E) staining and Masson trichrome staining of mouse heart after treatment. Quantification of fibrosis by Masson staining of collagen deposition on mice hearts. (**J-K**) Mouse cardiomyocytes after LNP treatment were stained with WGA and cardiomyocytes area were quantified by NIH Image J software. Mice were infused with 1000 ng/kg/min Ang II by osmic pump for 14days before vehicle or PDE3A2-modRNA treatment. Data were shown as mean ± SD. n = 6 mice for each group. *P* values were obtained by unpaired student *t*-test. \**p* < 0.05, \*\**p* < 0.01, \*\*\**p* < 0.001, \*\*\*\**p* < 0.0001.

### Single-nucleus RNAseq unraveled that PDE3A2-modRNA promoted physiological hypertrophy transcription of cardiomyocytes while inhibiting pathological hypertrophy-related signaling

To investigate the molecular mechanisms underlying the protective effects of PDE3A2-modRNA in PCH, we performed a single-nucleus RNAseq of mouse hearts in control (vehicle) or treated with PDE3A2-modRNA (**Figure 4** and **Online Figure 7**). Analysis of 21087 nuclei from the left ventricles of six hypertrophic mice (vehicle or PDE3A2-modRNA treatment) revealed significantmolecular alterations associated with the modRNA treatment. Using the Seurat package^31^, we clustered cells from all 6 samples into 15 distinct groups based on transcriptional similarities (**Figure 4A** and **Online Figure 7A**). Cell-type assignments, based on the top differentially expressed genes (**Online Figure 7B**), identified predominant populations of fibroblasts (15-25%), endothelial cells (∼36%), and cardiomyocytes (∼12.5%) (**Figure 4A-C**), consistent with prior studies^2^. Although PDE3A2-modRNA did not dramatically alter cell type proportions (**Figure 4B-C**), it substantially reshaped the transcriptional landscape across these cell types that induced total 2441 upregulated genes and 5017 downregulated genes from these cell types (**Figure 4D** and **Online Figure 7B**, **C**).

**Figure 4.**
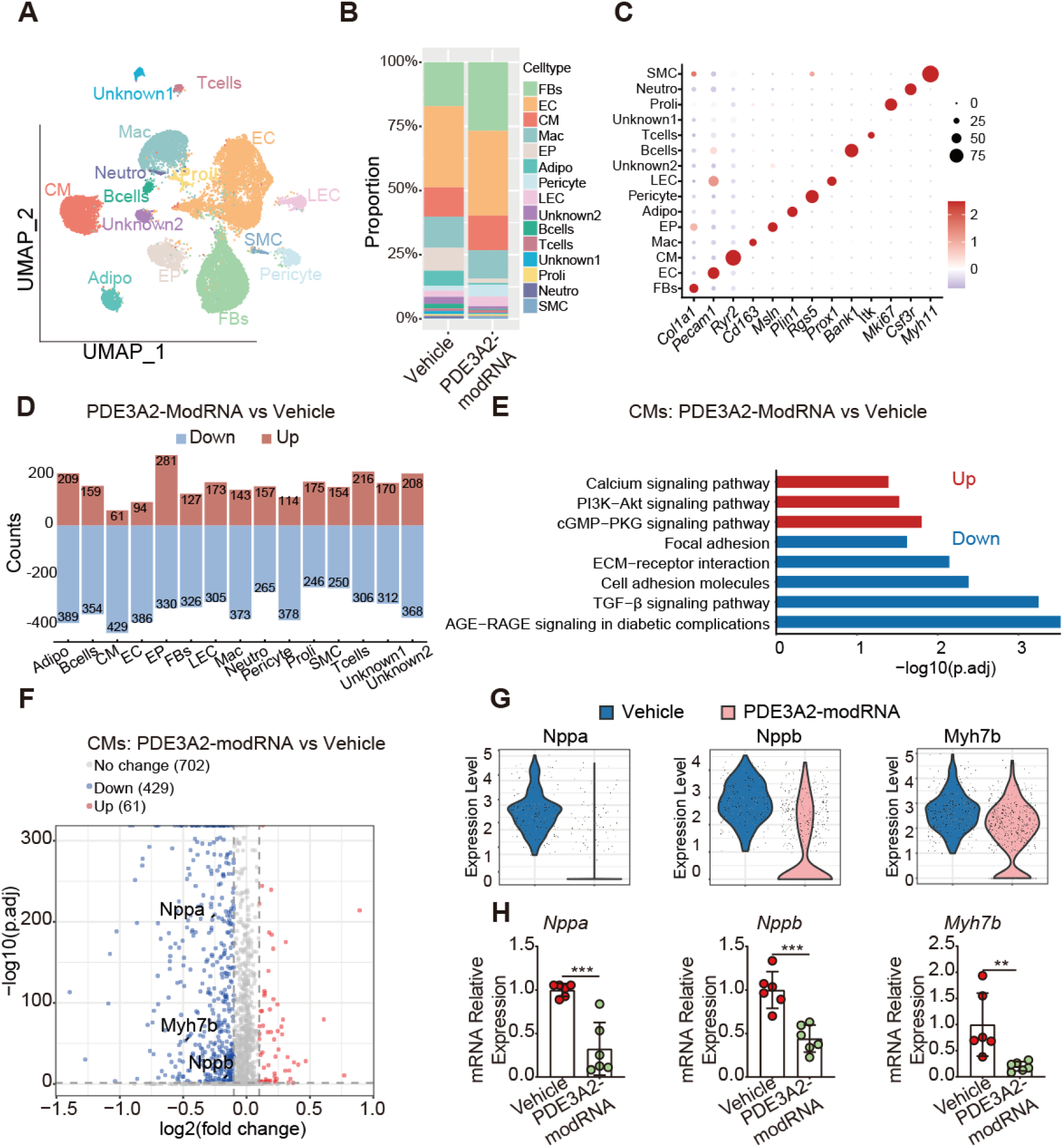
PDE3A2-modRNA alters hypertrophic mice heart transcription program and reduces cardiomyocyte fetal gene transcription. (**A**) Uniform Manifold Approximation and Projection (UMAP) showing single nuclei isolated from 3 Ctrl (Vehicle) and 3 PDE3A2 treated PCH mice left ventricles. (**B**) Cellular composition in the left ventricles from Vehicle and PDE3A2-modRNA treated hearts. (**C**) Dotplot showing expression of marker genes for each cluster. (**D**) The number of significant differentially expressed genes in distinct cell types by PDE3A2 vs Vehicle comparison. The red and blue color indicates the up-regulated or down-regulated genes, respectively, by PDE3A2-modRNA. Mice were *i.v.* injected with LNP control or LNP-encapsulated PDE3A2-modRNA (every 2 days; 20μg) after infusing with Ang II for 14 days. (**E**) GO analyses of DEGs for cardiomyocytes. Selected top categories are shown. (**F**) Volcano plot of differential expressed gene (DEG) in cardiomyocytes (CM) between Vehicle and PDE3A2-modRNA treatments. (**G**) The mRNA level of hypertrophic fetal genes (*Nppa*, *Nppb* and *Myh7b*) in individual CMs. (**H**) Quantitative mRNA assay of fetal genes in Vehicle and PDE3A2-modRNA treated mice hearts. For panel **H**, *P* values were obtained by unpaired student *t*-test. \*\**p* < 0.01, \*\*\**p* < 0.001.

Specifically in cardiomyocytes, PDE3A2-modRNA upregulated genes involved in the PI3K-Akt, cGMP-PKG signaling pathways (**Figure 4E**), as well as genes associated with Gap junctions-key regulators of cardiac adaptive responses and the development of physiological hypertrophy^32^. Activation of the PI3K-Akt and cGMP-PKG pathways is known to enhance physiological cardiomyocyte growth while suppressing pathological hypertrophy^33–35^. Meanwhile, PDE3A2-modRNA inhibited TGF-β signaling, ECM and cell adhesion pathways, which are key players that promote cardiac fibrosis (**Figure 4E**). PDE3A2-modRNA also down-regulated genes involved in pathological hypertrophy (**Figure 4F-G**), including *Nppa, Nppb, and Myh7b*, which are typically upregulated in pathological hypertrophy and downregulated in physiological hypetrophy^1^. qPCR analysis corroborated the downregulation of *Nppa, Nppb, and Myh7b* in mouse hearts induced by PDE3A2-modRNA (**Figure 4H**). Overall, these findings underscore the profound changes in transcriptional landscape of hypertrophic hearts, particularly cardiomyocytes, in response to PDE3A2-modRNA. Importantly, genes associated with physiological hypertrophy were up-regulated, while those involved in pathological hypertrophy were suppressed by the treatment, indicating that PDE3A2-modRNA may reverse the pathological cardiac hypertrophy.

### PDE3A2-modRNA reduced cardiomyocyte injury-responding functional trajectories, inhibiting inflammatory complement activation

To characterize the cardiomyocyte response following PDE3A2-modRNA treatment, we partitioned all cardiomyocytes into 9 subtypes (**Figure 5A-B**), according to previous study^2^. Notably, PDE3A2-modRNA altered the distribution of these cardiomyocyte subtypes (**Figure 5C**). Spearman correlation analysis unraveled that 9 CM subtypes could be grouped into 4 functional clusters (FCs) (**Figure 5D-E** and **Online Figure 8A-B**). Each FCs exhibited distinguished cardiomyocyte states, including differences in muscle contraction and molecular pathways (**Figure 5F-G**). Specifically, FC1 was the most abundant population (∼30% in vehicle group and ∼50% in PDE3A2-modRNA) and was characterized by the expression of *Ttn* (**Figure 5G** and **Online Figure 8A**), which generate passive force in sarcomere to regulate cardiac relaxation and it loss of function mutations are associated with hypertrophic cardiomyopathy^36,37^. FC3, which highly expressed fibrotic and wound-healing genes (**Figure 5F**), accounted for ∼50% of cardiomyocytes in PCH hearts (vehicle) and was reduced by PDE3A2-modRNA (**Figure 5G**). We identified FC2 and FC4 as injury-associated populations. FC2, marked by the expression of genes involved in cardiac hypertrophy response and vascular remodeling^2^, including *Gja1* and *Nppb*. FC4 was associated with wound healing involved inflammatory response and chemotaxis (**Figure 5F**), as marked by Cxcl12 and Cfd (**Online Figure 8A**). Further, pseudotime analysis using Slingshot revealed an injury-correlated trajectory that was initiated from FC1 towards FC3 (**Figure 5H**), which further bifurcated into FC2 (**Figure 5H**, Path 1) or FC4 (**Figure 5H**, Path 2). In addition, the greater fraction of FC1 following PDE3A2-modRNA treatment was attributable to the attenuated “differentiation” from FC1 to FC3 in Path 1 (**Online Figure 9A**). Overall, these results suggest PDE3A2 could rescue the contractile FC1 and align with the improved diastolic function in PCH mouse.

**Figure 5.**
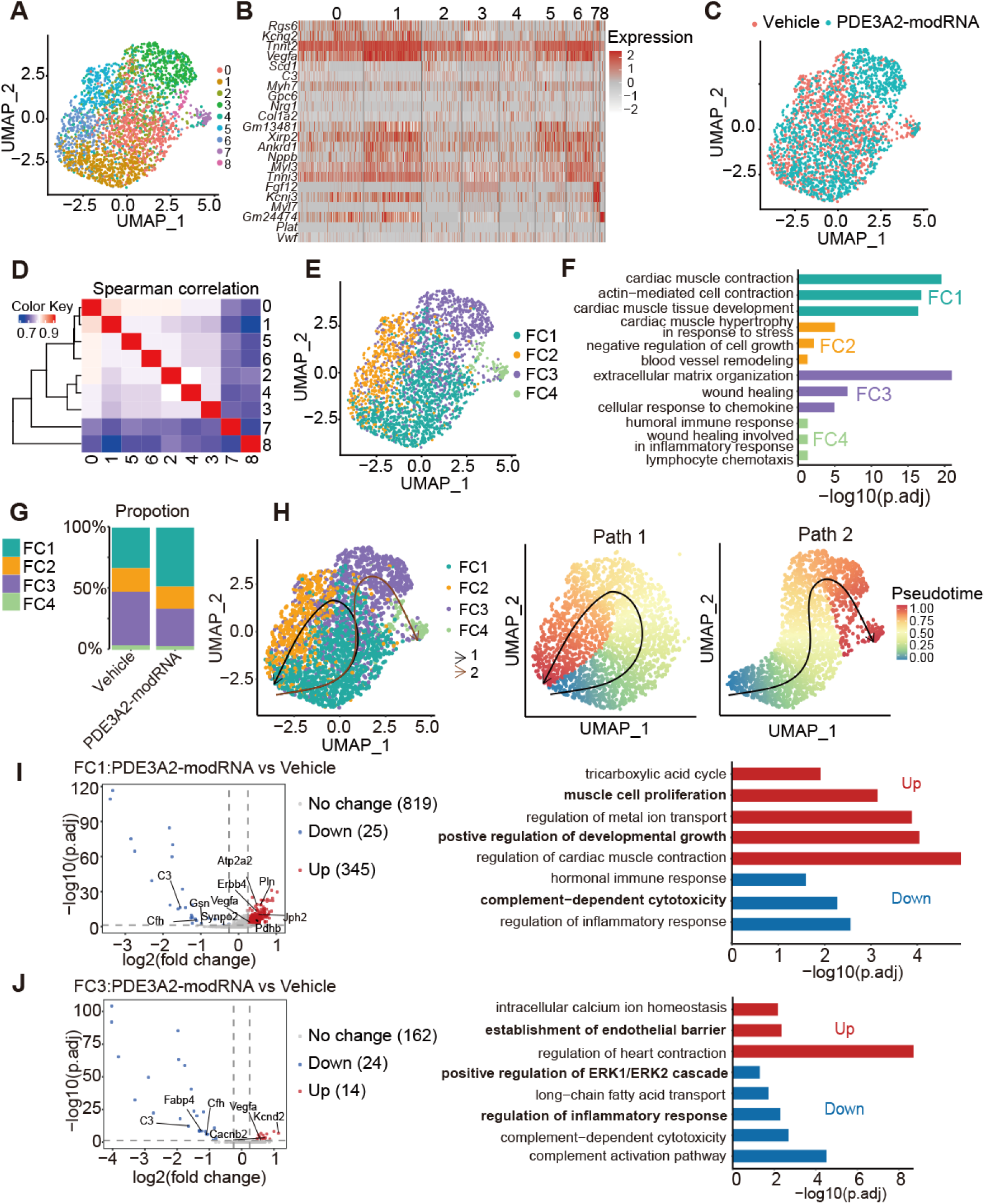
PDE3A2-modRNA modifies CM subclustered gene transcription profile in the PCH hearts. (**A**) UMAP showing CMs isolated from mouse hypertrophic hearts with LNP Ctrl or PDE3A2-modRNA after 14 days infusion of Ang II. CMs were marked by cluster numbers. CM [0-8]. (**B**) Heatmap depicting DEGs among cardiomyocyte (CM) clusters. (**C**) UMAP projection showing CMs from Vehicle and PDE3A2-modRNA treated mouse hypertrophic hearts. (**D-E**) 9 CM subtypes were grouped into 4 functional clusters (FCs) according to transcriptome similarity. Individual FCs were depicted in different colors. (**F**) Gene ontology (GO) analysis of specifically expressed genes in each FC. (**G**) Proportion of the 4 different FCs in Vehicle and PDE3A2-modRNA treated hearts. (**H**) Slingshot analyses showing CMs trajectory in pseudotime. Each color represents a FC. (**I-J**) Comparison of differential expressed genes (left) and pathways (right) between FC1 and FC3 in PDE3A2-modRNA relative to Vehicle group.

Our subsequent analysis was aimed to examine the transcriptomes of FC1 and FC3 affected by PDE3A2-modRNA treatment. We identified significant upregulation of (in total 345) genes in FC1 that were associated with cell proliferation and growth (*Erbb4*), TCA metabolism (*Pdhb*), and muscle contraction (*Atp2a2*, *Pln* and *Jph2*; **Figure 5I**). These gene signatures are typically persevered in physiological hypertrophy, offering protection against pathological hypertrophy^1,38,39^. Additionally, PDE3A2-modRNA downregulated genes in FC1 that are involved in inflammatory responses and complement-dependent cytotoxicity, namely C3 and Cfh (**Figure 5I**). Specifically, complement C3 activation is known to promote release of inflammatory cytokines and contribute to cardiac remodeling^40^. In the injury-responding FC3, PDE3A2-modRNA upregulated Vegfa-mediated angiogenesis while suppressing C3 activation and the ERK pathway (**Figure 5J**). Indeed, our result unraveled that PDE3A2-modRNA reduced the expression of *C3d* (marker of C3 activation; **Online Figure 10A**) while enhancing the excitation-contraction coupling-associated genes (*Pln, Atp2a2*, and *Jph2*; **Online Figure 10B-D**) and Vegfa (**Online Figure 11A**). Collectively, these findings highlight the impacts of PDE3A2-modRNA on the injury-associated cardiomyocyte trajectory, attenuating the transition from physiological FC1 to remodeling-associated FC3.

### PDE3A2-modRNA improved cardiac microenvironment by attenuating oxidative stress and immune response in non-cardiomyocytes

Imbalance energy metabolism, reactive oxidative stress and immune response promotes pathological deterioration of cardiac hypertrophy that involves various non-cardiomyocyte cell types, including fibroblasts, macrophages and adipocytes^41–43^. Among them, the activation of fibroblast into myofibroblast, characterized by extracellular matrix (ECM) secretion, promotes tissue stiffness and facilitates cardiac fibrosis. PDE3A2-modRNA treatment alleviated the transition of fibroblast into myofibroblast by reducing collagens secretion (**Online Figure 11B**) and α-SMA production (**Figure 6A-C**, **Online Figure 11C** and **Online Figure 12A**), as evidenced by the lower expression of myofibroblast marker genes, including *Col14a1*, *Col1a1*, *Col16a2* and *Acta2* (**Figure 6A**). Consistently, GO analysis confirmed that ECM, response to oxidative stress and adhesion-related signaling pathways were downregulated, whereas cardioprotective ATP biosynthesis process and endothelial cell development, were significantly up-regulated by PDE3A2-modRNA (**Figure 6B**). Accumulating evidence suggests inflammation as a key regulator in the initiation and progression of cardiac diseases, including PCH^44^. In hypertrophic hearts, macrophage polarization into M1 to produce pro-inflammatory cytokines, like IL-1β, IL-6 and CCL4 (a ligand for CCR5), and thus facilitates myofibroblast activation and cardiac fibrosis^45^. We found that PDE3A2-modRNA significantly suppressed the macrophage pro-inflammatory TLR4 signaling and oxidative stress, which were drivers of M1 macrophage polarization. And PDE3A2-modRNA down-regulated inflammatory cytokines (IL-1β and IL-6) and chemokines (**Figure 6D-E** and **Online Figure 13**). On the other hand, M1 to M2 macrophage switch improves cardiac repair by enhancing efferocytosis and phagocytosis of apoptotic and dead cardiomyocytes^46^, thereby reducing inflammation and attenuating heart damage. We found PDE3A2-modRNA shifts the pro-inflammatory M1 macrophages towards cardioprotective M2 macrophages, as it elevated M2 macrophage marker, CD163, which was confirmed with mouse heart immunostaining (**Figure 6D-F** and **Online Figure 12B**). Adipocytes play a critical role in the endocrine regulation of cardiac microenvironment by secreting a variety of metabolites and thus by communicating with other cell types, such as fibroblast, macrophages and cardiomyocytes (**Figure 6G-I**)^47,48^. Of note, deficient energy metabolism in adipocytes generates large amount of reactive oxygen species and induces oxidative stress to local microenvironment^49^. We found PDE3A2-modRNA enhanced fatty acid metabolism and thermogenesis as it promoted expression of Adrb3, Ppargc1α, Acly and Fabp4 in adipocytes (**Figure 6G-H** and **Online Figure 12C**). These genes have been shown to promote lipolysis and protect against cardiac damage^43,50,51^. This sequencing result was confirmed by elevation of expression of β_3_-AR by PDE3A2-modRNA (**Figure 6I**). Meanwhile, PDE3A2-modRNA treatment reduced interleukin production and hydrogen peroxide catabolic process, which are consistent with the downregulation of oxidative stress and inflammation in fibroblasts and macrophages. Collectively, the single-cell RNA sequencing analysis indicated underrecognized effects of PDE3A to rebuild cardiac microenvironment by reducing the oxidative stress and immune response.

**Figure 6.**
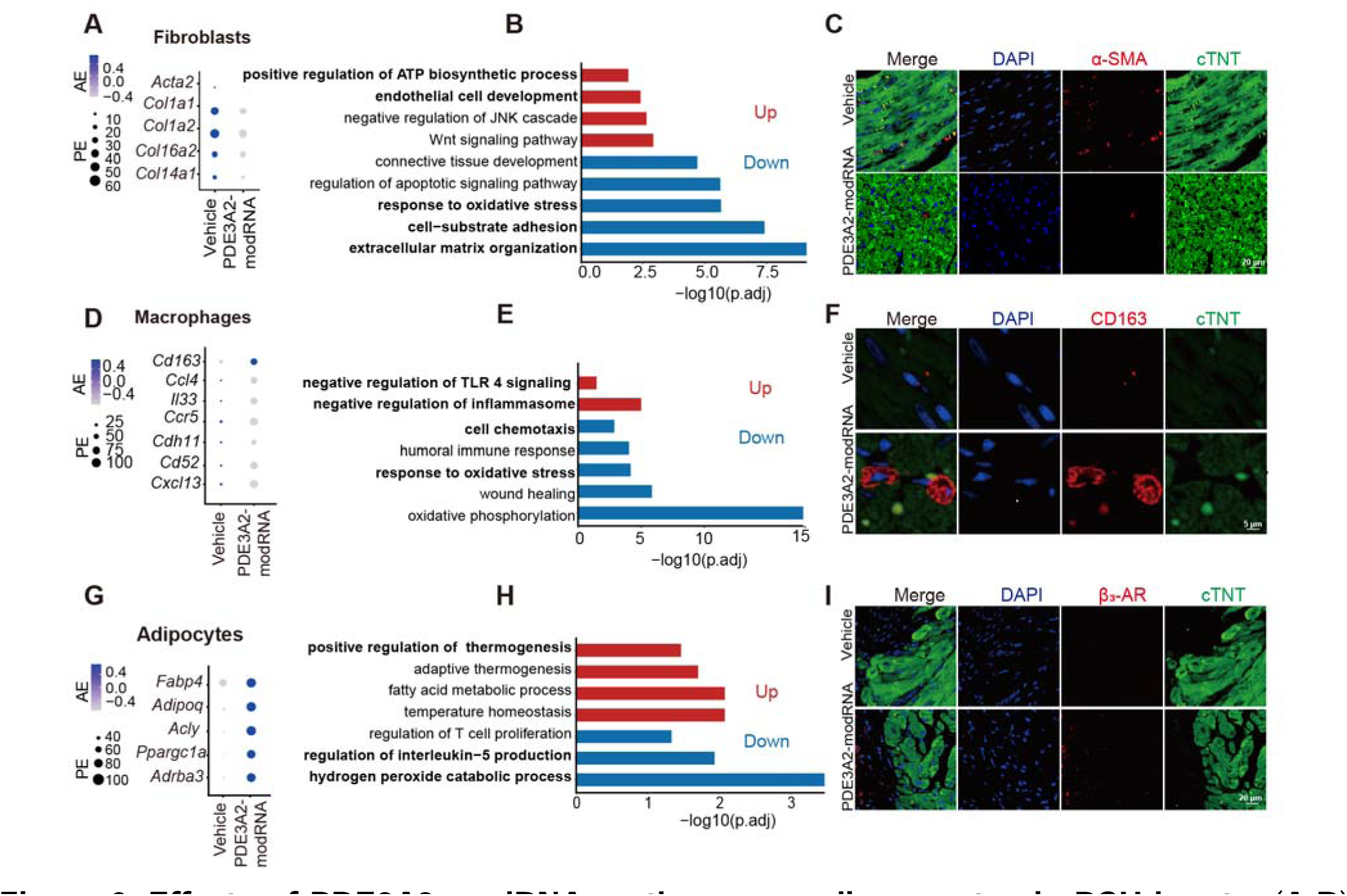
Effects of PDE3A2-modRNA on the non-cardiomyocytes in PCH hearts. (**A-B**) Typical genes and pathways altered by PDE3A2-modRNA in fibroblasts. (**C**) α-SMA expression in PCH hearts treated with Vehicle or PDE3A2-modRNA. (**D-E**) Typical genes and pathways altered by PDE3A2-modRNA in macrophages. (**F**) PDE3A2-modRNA promoted M2 macrophage marker expression (CD163) in PCH hearts. (**G-H**) Typical genes and pathways altered by PDE3A2-modRNA in adipocytes. (**I**) PDE3A2-modRNA elevated β_3_-adrenergic receptors in adipocytes.

## Discussion

PCH is a common precursor to heart failure, a major cardiovascular disease with increasing prevalence and mortality worldwide^2^. Using subcellular-localized FRET biosensors, we identified heterogeneous changes in cAMP microdomains in PCH, specifically finding hyperactive nuclear cAMP linked to reduced PDE3A2 levels. ModRNA targeting PDE3A2 restored nuclear PDE3A2 levels, reduced NLS-cAMP without affecting cytosolic cAMP, and improved cardiac function *in vivo*. Single-nucleus transcriptomics revealed that PDE3A2-modRNA promoted gene programs associated with physiological hypertrophy in cardiomyocytes, while suppressing pathological hypertrophy-related signaling pathways. Specifically, PDE3A2-modRNA reduced remodeling-associated cardiomyocyte functional subclusters by attenuating injury-responding functional trajectories and inhibiting inflammatory complement activation. Additionally, PDE3A2-modRNA treatment rebuilds the cardiac microenvironment by modulating fibroblasts, macrophages, and adipocytes, leading to reduce oxidative stress and immune response. PDE3A2-modRNA presents a therapeutic strategy to reverse pathological hypertrophy by targeting nuclear cAMP microdomain.

### Translation perspective of targeting compartmentalized cAMP to treat cardiac diseases

Subcellular cAMP is organized at the nanometer scale^8,52–54^, and targeting the cAMP microdomains is emerging as a potential therapeutic strategy due to its high fidelity and selectivity within specific cellular compartments^9,12^. Here, we characterized cAMP microdomains in PCH mice model and unmasked heterogeneous alterations of compartmentalized cAMP signaling using subcellular-localized PKA biosensors. While PKA response at PM was diminished, nuclear PKA was hyperactive in PCH. This heterogeneity of cAMP-PKA signaling is consistent with a previous study showing reduced PKA activity at the sarcolemma in hypertrophic rabbit myocytes^55^, suggesting that the PCH-induced differential modifications to cAMP microdomains are conserved across these species. Reduced cAMP-PKA activity at sarcolemma can impair PKA-dependent phosphorylation of multiple ion channels, ultimately leading to impaired cardiac contraction^56^. The distinct alterations in these two local cAMP microdomains may be attributed to the desensitization and internalization of adrenergic receptors in PCH^12,57^, though the underlying molecular mechanisms require further investigation. While PDE3A2 was recently demonstrated to be located to nucleus, PDE3A1 is localized to intracellular membranes^58^ to associate with sarcoplasmic/endoplasmic reticulum Ca^2+^-ATPase and critically modulate cardiomyocyte Ca^2+^ cycling and cardiac systolic function^14,59^. Understanding the intricate cAMP landscape within the heart and unraveling the mechanisms governing compartmentalized signals are essential for translating these insights into effective clinical interventions.

NLS-cAMP has recently been shown to modulate gene expression^60^ and PKA-dependent phosphorylation of histones^22,61,62^, which are involved hypertrophic gene transcription in neonatal cardiomyocytes. A recent study demonstrated that PDE3A2 interacts with histone deacetylase 1 (HDAC1) in neonatal cardiomyocytes. Deficiency of PDE3A2 promoted PKA-dependent phosphorylation of HDAC1, leading to increased hypertrophic gene transcription *in vitro*. We observed a hyperactive nuclear cAMP in PCH mouse hearts, due to reduced PDE3A2 but not PDE3A1. This finding may explain why non-selective PDE3 inhibitors can transiently improve cardiac systolic function, but are associated with increased mortality ^63^. Targeting the specific PDE3A2 isoform, rather than PDE3 globally, avoids interference with cytosolic or membrane-bound cAMP, offering a promising strategy to fine-tune dysregulated gene transcription in cardiac hypertrophy with minimal impacts on physiological cardiac function, including electrophysiology.

Dissecting the downstream signaling regulated by NLS-cAMP is essential for understanding and identification of therapeutic strategies targeting NLS-cAMP. Our transcriptomic analysis together with immunostaining reveals a critical role of PDE3A2-mediated suppression of NLS-cAMP in enhancing myocyte contraction and reducing inflammation in PCH. Specifically, PDE3A2 increased titin-positive cardiomyocytes and upregulated genes such as *Atp2a2, Pln* and *Jph2*, which are the key for improving calcium handling and enhancing diastolic function in PCH hearts^64,65^. Meanwhile, PDE3A2 downregulated *C3* and *Cfh*, thereby inhibiting complement activation, a major regulator for the pathological microenvironment of a heart^40^. In addition, PDE3A2-mediated regulation of NLS-cAMP synergistically reduced oxidative stress and inflammation in fibroblasts, macrophages, and adipocytes, contributing to the rebuild of PCH cardiac microenvironment. Interestingly, while the multifunctional role of cAMP in these non-cardiomyocytes is recognized, the compartmentalization of cAMP in these cells remains largely unexplored^61^. Our findings suggest that the therapeutic effects of PDE3A2 in PCH extend beyond cardiomyocytes to non-cardiomyocytes. This study highlights the pivotal role of PDE3A2 in modulating contractile and inflammatory gene expression, providing novel insights into the molecular mechanisms underlying the PDE3A2 regulation of physio(patho)ology.

### ModRNA as a feasible strategy for isozyme-specific PDE targeting in the heart

Targeting a specific isoform or variant of PDEs remains a major challenge with conventional pharmacological. Modified mRNA (modRNA) emerges as a potential strategy that can drive specific PDE variant expression. ModRNA-dependent gene expression has been clinically proved to be efficient, titratable, and minimally immunogenic with bell-shaped pharmacokinetics and no risk of genomic integration^19^. In fact, synthetic modRNA has shown success in both diseased animal models and human hearts^66,67^. For example, modRNA encoding VEGF-A has advanced to Phase II clinical trials for treatment of ischemic hearts. Our study, for the first time, demonstrates that modRNA can be effective in isozyme-specific interventions for managing chronic cardiac diseases, such as hypertrophy. Additionally, although modRNA is typically administered to the heart via local cardiac injection, we found that *i.v.* injection of modRNA is also effective, achieving approximately 40-50% efficiency compared to intramyocardial injection, making it a more feasible approach in this work. Surprisingly, despite the transient gene expression induced by modRNA, which typically lasted 8-12 days, three doses of modRNA significantly alleviated PCH. This suggests that *i.v.* administration of modRNA may provide a viable treatment option for cardiac diseases *in vivo*.

Future studies should focus on developing cardiomyocyte-targeted modRNA tools for clinical applications. With its transient and controllable target-gene expression, modRNA technology offers an efficient and safe gene therapy approach, holding potential for facilitating effective treatments for numerous cardiovascular diseases by enabling isozyme-specific interventions that selectively target subcellularly biased signaling microdomains.

### Limitations of the study

Utilizing a variety of molecular and cellular experiments, we demonstrated that PDE3A2-modRNA alleviates PCH in mice. However, our preventive treatment strategy (administering modRNA concurrently with inducing PCH) does not mirror the clinical scenario, where treatment begins after diagnosis. Future studies should examine the effects of PDE3A2-modRNA in a therapeutic regimen, whereby the treatment started until the cardiac dysfunction and fibrosis are developed. Furthermore, our findings are based on an Ang II-induced mouse model. It remains unclear whether PDE3A2-modRNA would be effective in other models of PCH, such as pressure overload induced by transverse aortic constriction surgery or in the context of human HCM. Finally, although PDE3A2-modRNA treatment shifted the hypertrophic transcriptome, the precise molecular mechanisms driving this beneficial transition require further elucidation.

## Methods

### Sex as a biological variable

Our study exclusively examined male mice. It is unknown whether the findings are relevant for female mice.

### Reagents

Angiotensin II (Ang II, A800620) was obtained from Macklin (Shanghai, China). Norepinephrine (NE, HY-1315A), Forskolin (FSK, HY-15371) and 3-Isobutyl-1-Methylxanthine (IBMX, HY-12318) were purchased from MedChemExpress (Shanghai, China).

### Animals and experiment design

Male wild-type C57BL/6J mice, aged 8 weeks, were purchased from Gempharmatech Co.Ltd (Guangdong, China). All mice underwent a one-week adaptive feeding before experiments. The animal experiments were approved by the Institutional Animal Care and Use Committees (IACUC) of Southern University of Science and Technology (Protocol No. SUSTech-JY202306004-202404A1). Mice were maintained in a specific-pathogen-free (SPF) environment with a 12-hour light/dark cycle and a controlled temperature of 22-24 °C.

For the Ang II-induced cardiac pathological hypertrophy (PCH) model, all wild-type C57BL/6J mice were injected with Ang II (1000 ng/kg/min) for 14 days using osmotic minipumps, as previously described^17^. Briefly, mice were anesthetized with isoflurane (5% for induction and 2%-3% for maintenance) in pure oxygen flow (0.8 L/min). Ventilation was applied at a respiratory rate of 110 to 130 breaths/min and a tidal volume of 0.2 mL for the minipumps implanting. Analgesia (buprenorphine 0.1 mg/kg) was given immediately after surgery and after recovery.

14 Days post-implantation of the pumps, the mice were randomly divided into PCH-Control and PDE3A2-modRNA treated groups. LNP encapsulated PDE3A2-modRNA (20 μg) was administered by tail vein injection once every two days, for a total of three injections.

### Design and synthesis nucleic acid drug of PDE3A2

PDE3A2 protein sequences were retrieved from NCBI (Uniport: UP000314294). To facilitate secretion, the leader sequence of tissue plasminogen activator (TPA) was added at the N-terminus, and a 6×His tag was appended at the C-terminus of the construct. For further optimization of the sequence, 5’ and 3’ UTRs (human beta globulin UTR) were added as previously described^18^. Additionally, KOZAK sequence was introduced at the 5’ end of the open reading frame (ORF) and a polyadenylation (PolyA) sequence at the 3’ end. These gene fragments were synthesized and cloned into the pVAX1 vector to obtain template plasmids pVAX1-PDE3A2.

### In vitro transcription and purification of mRNA

In vitro mRNA synthesis was conducted as previously described^18^. Briefly, DNA transcription was mediated by T7 or SP6 polymerase with the promoter sequence– ATTTAGGTGACACTATAG. The steps of *in vitro* transcription are as follows: (1) mix RNase free water and NTPs; (2) Add CleanCap® Reagent AG (3’ OMe), mix, and centrifuged to collect the supernatant; (3) Add 10×Transcription Buffer, mix, and centrifuge to collect the supernatant; (4) Add the DNA template; (5) Add RNase Inhibitor, Yeast Inorganic Pyrophosphatase, and T7 RNA Polymerase, mix well, and centrifuge to collect the supernatant; (6) Incubate the reaction at 37 °C for 2-3 hours; (7) Add DNAaseI to remove the template DNA, then purify the transcript using Monarch RNA cleanup kit (Thermo Fisher Scientific, USA). The concentration of mRNA was measured using a NanoDrop 2000c UV spectrophotometer, and the purity of the mRNA was determined by capillary electrophoresis 2100 Bioanalyzer.

### Determination of particle size, uniformity, and encapsulation efficiency of lipid micro particle (LNP)-encapsulated modRNA

LNPs were prepared by dissolving ionizable lipid (DLin-MC3-DMA), cholesterol, DSPC, and polyethylene glycol in ethanol (ratio 50:38.5:10:1.5) to a concentration of 10 mg/ml, forming the organic phase. The mRNA was diluted in sodium citrate buffer (50 mM, pH 4) to 0.1 mg/mL, forming the aqueous phase. These phases were mixed (1:3) using a microfluidic device at 12 ml/min, yielding the LNP-mRNA mixture. The mixture was diluted 40-fold with PBS and centrifuged using an Amicon® Ultra-15 filter at 4000 × g for 30 minutes at 4 °C, repeating three times. The final product of LNP-encapsulated mRNA was stored at 4 °C.

Particle size and uniformity were measured by dynamic light scattering with HORIBA-SZ100 equipment. Encapsulation efficiency was determined using the Quant-iT™ RiboGreen™ RNA kit, measuring free nucleic acid before and after 1% Triton-100 treatment.

### modRNA-LNPs cytotoxicity assay

HEK 293A cells (purchased from ATCC) were cultured adherently, and different doses of modRNA were added to assess cytotoxicity. Briefly, 4-6 hours after seeding cells in 96-well plates, LNP-modRNA at different doses: 1 µg, 5 µg and 20 µg were added. After 24 hours, 10 µl of CCK8 (Vazyme, Nanjing, China) was added to each well and cultured for 0.5 to 4 hours. Cell viability was quantified by measuring the OD at 450 nm using a microplate reader.

### Administration of modRNA *in vivo*

ModRNA was delivered to mice via intramyocardial, or intravenous injection, as previously reported^19, 20^. For intramyocardial injection, 20 µg of modRNA in 20 µL was administered to six sites around the left ventricles of the mice^19^. For intravenous injection, 20 µg of modRNA in 20 µL was administered intravenously (*i.v.*) via the tail vein^20^.

### Echocardiography

Mice cardiac function was measured by ultrasound imaging system (Vevo1100, Visual Sonic, Canada) as previously described^21^. Briefly, mice were anesthetized with 1-2% isoflurane and imaged by M-model, tissue doppler, and color doppler mode were applied to evaluate the systolic and diastolic cardiac function.

### Adult Mice Cardiomyocytes Isolation

Adult mice cardiomyocytes isolation was performed as previously described^44^. Briefly, 3-5% isoflurane was used to anesthetize mice. And then, hearts were quickly excised and mounted on a Langendorff perfusion apparatus. A solution containing collagenase and protease (0.5 mg/mL collagenase and 0.1 mg/mL protease) were perfused into the hearts. Freshly isolated cardiomyocytes were cultured and set aside for upcoming experiments.

### Cell Culture

HEK 293A cells and Adult cardiomyocyte isolated from mouse were cultured in a 4.5 g/L glucose DMEM medium with 10% FBS and 1% penicillin/streptomycin in 5% CO_2_ at 37 °C as previously described^11^.

### Immunofluorescence staining

Paraffin-embedded heart tissue sections and cultured cells were used for immunofluorescence staining. Briefly, samples were dewaxed and deparaffinized with xylene. Sections were kept moist in water until antigen-antibody repair. The slices were boiled in sodium citrate buffer for 15 minutes for antigen retrieval, then washed with PBS three times (total 15 minutes). Sections were covered with blocking buffer for 30 minutes, then incubated with primary antibodies (PDE3A, 1:200, PU298267S, Abmart; α-SMA, 1:200, 19245, Cell Signaling Technology; CD163, 1:200, 16646-1-AP, Proteintech; β_3_-AR, 1:200, PS05942S, Abmart; cTnT, 1:200, 68300-1-Ig, Proteintech; Anti-His-tag, 1:200, AE086, ABclonal) overnight at 4 °C or for 1 hour, washed with PBS three times, and incubated with secondary antibodies (Goat Anti-Rabbit Alexa Fluor 594, 1:200, M21014M, Abmart; Goat Anti-Mouse Alexa Fluor 488, 1:200, M21011, Abmart) at room temperature. Finally, sections were washed with PBS (15 minutes, three times), counterstained with DAPI, and observed using a Nikon microscope.

### Western blotting

Western blot was conducted as previously reported^11^. In brief, a total of 30 μg protein were loaded to 10% SDS-PAGE gel electrophoresis and transferred onto 0.2 μm nitrocellulose filter membrane (HATF02500, Merck millipore). To reduce the amount of background produced by non-specific binding, NC membranes were blocked with 5% skim milk for 1 hour, at room temperature. And membranes were incubated with primary antibodies (PDE3A, 1:1000, PU298267S, Abmart, Anti-His-tag, 1:1000, AE086, ABclonal) at 4 °C for overnight, after washing the membranes, then by an incubation with anti-rabbit HRP-conjugated second antibody at room temperature for 1 hour. Protein blots were visualized by chemiluminescence detection through ECL (BL520B, LABGIC). Band intensities were quantified with Image J software (NIH, Bethesda, MD, USA).

### Fluorescent Resonance Energy Transfer (FRET) Assay

FRET-based PKA biosensors were expressed in 293A cells and isolated cardiomyocytes before FRET measurement using Nikon TiE2 microscope. Subcellular-localized FRET biosensors (NLS-AKAR3, NES-AKAR3 and PM-AKAR3) have been reported previously^22^. FRET images of YFP/CFP were acquired as previously described and analyzed. ΔYFP/CFP ratio was normalized to baseline, and its changes reflect the PKA activity response, as previously described^22^.

### Nuclei isolation sorting from mouse hearts and single nucleus RNA-sequencing library preparation

Mouse heart tissues were harvested and washed in pre-cooled PBSE (PBS buffer containing 2 mM EGTA). Nuclei isolation was performed using GEXSCOPE® Nucleus Separation Solution (Singleron Biotechnologies, Nanjing, China) according to the manufacturer’s instruction. The isolated nuclei were resuspended in PBSE to a concentration of 10^6^ nuclei per 400 μL, filtered through a 40 μm cell strainer, and counted with Trypan blue. Nuclei enriched in PBSE were stained with DAPI (1:1,000) (ThermoFisher Scientifc, D1306). Nuclei were defined as DAPI-positive singlets.

The concentration of the single nucleus suspension was adjusted to 3-4 × 10^5^ nuclei/mL in PBS. The single nucleus suspension was then loaded onto a microfluidic chip (GEXSCOPE® Single NucleusRNA-seq Kit, Singleron Biotechnologies) and snRNA-seq libraries were constructed according to the manufacturer’s instructions (Singleron Biotechnologies). The resulting snRNA-seq libraries were sequenced on an Illumina novaseq 6000 instrument with 150 bp paired-end reads.

### Processing of snRNA-seq data

Raw reads from snRNA-seq were processed to generate gene expression matrixes using the CeleScope pipeline (v1.9.0, https://github.com/singleron-RD/CeleScope). Briefly, raw reads were first processed with CeleScope, where low-quality reads were removed, and Cutadapt (v1.17) was used to trim poly-A tail and adapter sequences^23^. Cell barcodes and UMI were extracted. Next, STAR (v2.6.1a^24^) to map the reads to the reference genome GRCm38 (ensembl version 92 annotation). UMI counts and gene counts of each cell were obtained using featureCounts (v2.0.1)^25^, generating expression matrix files for subsequent analysis.

For each snRNA-seq dataset, quality control steps included removing low-quality cells and filtering by mitochondrial content using the Seurat R package^62^. The cell passing the min.feature 200 and gene satisfying min.cells 3 were kept for downstream analysis. Cells with over 10% mitochondrial content were removed. Furthermore, to check the quality and remove any multiplets (or double cells) of the single-cell data, we performed Seurat-based filtering of cells based on nFeature_RNA (200 to 2,500). After filtering, 11418 and 9669 cells were retained for downstream analysis in the Ang II group and PDE3A2-modRNA group, respectively.

### Integrative analysis of snRNA-seq datasets

The integration methods (Seurat v4) were applied for the downstream analysis using 20 principal compotents (PCs=20) and a resolution=0.4. Specifically, integrative analyses were performed following the Seurat pipeline (https://satijalab.org/seurat/articles/integration_introduction.html). For the Seurat pipeline (V.4.2.0), we followed the routine integration process. Log-normalization was carried out using the NormalizeData function; selection of highly variable genes was performed using the FindVariableFeatures function (VST method); correction of batch effects across samples was performed using the FindIntegrationAnchors and IntegrateData functions; dimensionality reduction was performed using the RunPCA and RunUMAP functions; neighbour network construction was performed using the FindNeighbors function. The marker genes for each population were identified using the FindMarkers function with the Wilcoxon rank-sum test for all pipelines.

### Differential expression analysis between Vehicle and PDE3A2-modRNA

We identified differentially expressed genes between vehicle and PDE3A2-modRNA-treated group in each common cell population. Briefly, we used the FindMarkers function with the Wilcoxon rank-sum test for all pipelines. A gene was considered differentially expressed if |log2FC|>1 and the adjusted p<0.05 (the multiple testing correction is the Benjamini and Hochberg) as a differentially expressed gene. We then performed pairwise comparisons of transcriptomes between PDE3A2-modRNA and Vehicle-treated samples (FC1: PDE3A2-modRNA vs Vehicle, FC3: PDE3A2-modRNA vs Vehicle) using the FindMarkers function. Genes with p_val_adj < 0.05 and |avg_log2FC|>0.25 were used to generate a consensus list of differentially expressed genes.

### Pathway enrichment analysis

The Kyoto Encyclopedia of Genes and Genomes (KEGG) and Gene Ontology (GO) analysis were performed using ClusterProfiler (V.3.18.1), a R package in Bioconductor. Pathways with p_adj value less than 0.05 were considered as significantly enriched. Gene Ontology gene sets only including biological process categories were used as reference.

### Subpopulation analysis of vehicle and PDE3A2-modRNA groups

We performed the subclustering analysis of cardiomyocytes (CM) populations. Specifically, dimensionality reduction was applied using RunPCA (k=10), followed by clustering with the FindCluster (Louvain algorithm, resolution=0.8) to identify the CM subpopulations.

### CM Trajectory inference analysis

Inference of pseudo-temporal trajectories within the CM population was conducted using Slingshot R package version 2.12.0. Slingshot trajectory was computed based on the UMAP dimensionality. The inferred trajectory was then visualized on CM subcluster UMAP as an arrowed curve. Trajectory-coupled pseudotime was inferred and displayed on the UMAP after removing the cells with NA pseudotime values.

## Statistical Analysis

All data were presented as mean ± standard deviation (SD). Group sizes were set by a priori power analysis for a two-tailed, two-sample t-test (α=0.05, power=0.8) to detect a 10% endpoint difference. Mice were grouped with blinding and randomization during the experiments. Fully blinded analysis was performed by different persons carrying out the experiments and analysis, respectively. No experimental animals or data were excluded from the analysis. Representative figures and images reflected each experiment’s average level.

Normality of the data was assessed using the Shapiro-Wilk test in GraphPad Prism 8 with significance at alpha = 0.05 (GraphPad Inc., San Diego, CA). The unpaired Student *t*-test was utilized for comparisons of two groups. One-way ANOVA was performed for comparison among three groups or > 3 groups followed by Tukey’s post-hoc test. A *P* value < 0.05 was considered statistically significant.

## Study approval

All animal studies were approved by Southern University of Science and Technology’s IACUC (Approval No. SUSTech-JY202306004-202404A1).

## Data availability

Sequencing data are available from the corresponding author upon reasonable request.

### Authors contributions

YW and HN conceived and designed the research; WX, ML, QG, and XS performed experiments and data analysis; YW, HN, WX and ML drafted the manuscript; YW, HN, WX, ML, QG, SL, XYK, GQ and HL conducted a critical revision of the manuscript; HN and XS provided important experimental resources. All authors revised the manuscript and approved the final manuscript. The co-first authors, WX, ML and QG, contributed equally to the data generation, analysis and manuscript writing.

### Funding support

This work was supported by National Natural Science Foundation of China, Grant/Award Numbers: 82470293 and 32300948.

## Acknowledgements

We thank Professor. Peng Wang and XY. Wang, Southern University of Science and Technology for providing modRNA and its-related technical support.

